# Biallelic mutations in cancer genomes reveal local mutational determinants

**DOI:** 10.1101/2021.03.29.437407

**Authors:** Jonas Demeulemeester, Stefan C. Dentro, Moritz Gerstung, Peter Van Loo

## Abstract

The infinite sites model of molecular evolution requires that every position in the genome is mutated at most once^1^. It is a cornerstone of tumour phylogenetic analysis^2^, and is often implied when calling, phasing and interpreting variants^3,4^ or studying the mutational landscape as a whole^5^. Here we identify 20,555 biallelic mutations, where the same base is mutated independently on both parental copies, in 722 (26.0%) bulk sequencing samples from the Pan-Cancer Analysis of Whole Genomes study (PCAWG). Biallelic mutations reveal UV damage hotspots at ETS and NFAT binding sites, and hypermutable motifs in *POLE*-mutant and other cancers. We formulate recommendations for variant calling and provide frameworks to model and detect biallelic mutations. These results highlight the need for accurate models of mutation rates and tumour evolution, as well as their inference from sequencing data.

---

Recent studies have shown systematic variation in mutation rates across the genome, resulting in specific hotspots^5–7^. In addition, breakdown of the infinite sites assumption at the scale of individual single nucleotide variants (SNVs) was flagged up in single cell tumour sequencing data as a potential confounder during phylogenetic reconstruction^8^. It is unclear however, whether mutational recurrence is likely to be observed in practice within bulk tumour samples. Population averaging and limited long-range information carried by short-read bulk sequencing make it difficult to directly assess the validity of the infinite sites model.

During the evolution of a single diploid lineage, four classes of infinite sites model violations may be considered (**Figure 1**): (i) biallelic parallel, where two alleles independently mutate to the same alternate base; (ii) biallelic divergent, by independent mutation of two alleles, each to another base; (iii) monoallelic forward, where one variant is mutated to another; and (iv) monoallelic back, whereby an earlier variant reverts back to wild type. We focus on biallelic mutations – which can also serve as a proxy for parallel events in different lineages – hypothesising these may be observed directly in bulk tumour genome sequencing data. Loss of a variant owing to large-scale genomic deletion is not considered, as it does not strictly contradict the infinite sites assumption, yet such events should be adequately assessed when interpreting cancer genomes^2,8,9^.

**Figure 1.**
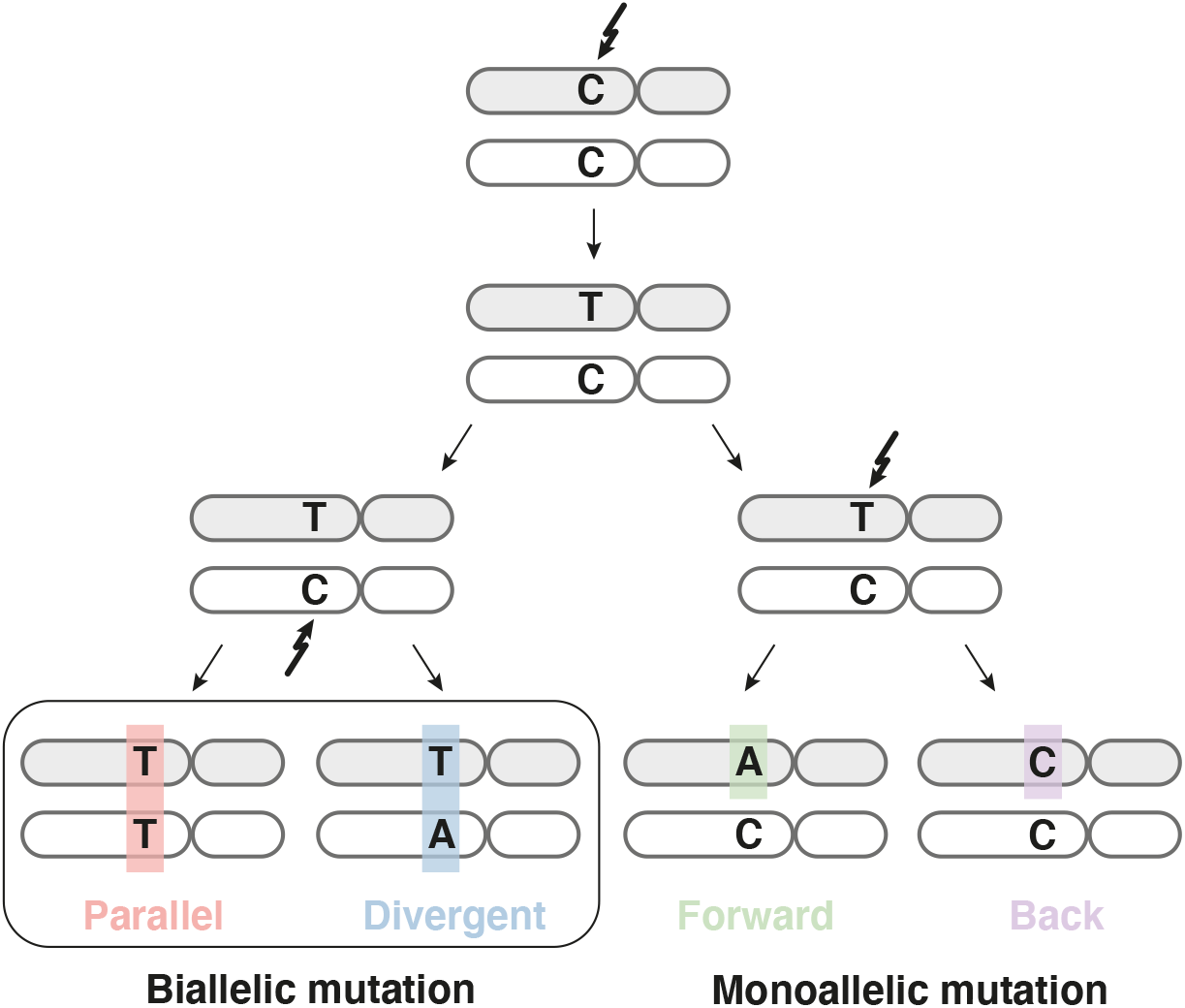
Possible violations of the infinite sites assumption in a single clonal lineage. Two subsequent mutations at a diploid locus can affect the same or alternate alleles. Depending on the base changes, there are four scenarios: biallelic parallel or divergent mutations affect separate alleles, whereas monoallelic forward and back mutation hit the same allele twice.

To assess the landscape of infinite sites violations, we start with a simulation approach using the PCAWG dataset of 2,658 whole-genome sequenced cancers. We resample a tumour’s observed mutations, preserving mutational signature exposures (96 mutation types, trinucleotide contexts)^10,11^ but otherwise assuming uniform activity of mutational processes across the callable regions of a diploid genome (uniform permutation model; **Table S1**, **Methods**). Given that mutation rates are most certainly not uniform and that any deviation from uniformity will only increase the number of violations^5^, this derives a lower bound. Even at this lower bound, these simulations indicate at least one violation in 147 tumours (5.5%, **Figure 2a**). Overall, biallelic parallel mutations represent the most common class of infinite sites model violation, with different tumour types showing different contributions from the other classes. A second simulation approach, resampling (without replacement) mutations from tumours of the same cancer type with similar mutational signature activities, confirms these observations (neighbour resampling model; **Figure 2b, Table S2**, **Methods**). Consistent differences between the simulators, in the number of violations per tumour type, inform on the non-uniformity of the mutational processes, *i.e.* a reduced “effective genome size” (akin to the population genetics concept of effective population size). With a median 75-fold excess violations compared to the uniform permutation model, the effective genome size is smallest in Lymph-BNHL (~2,782/75 = 37Mb ; **Figure 2c, Methods**), driven by somatic hypermutation recurrently targeting specific regions^12^.

**Figure 2.**
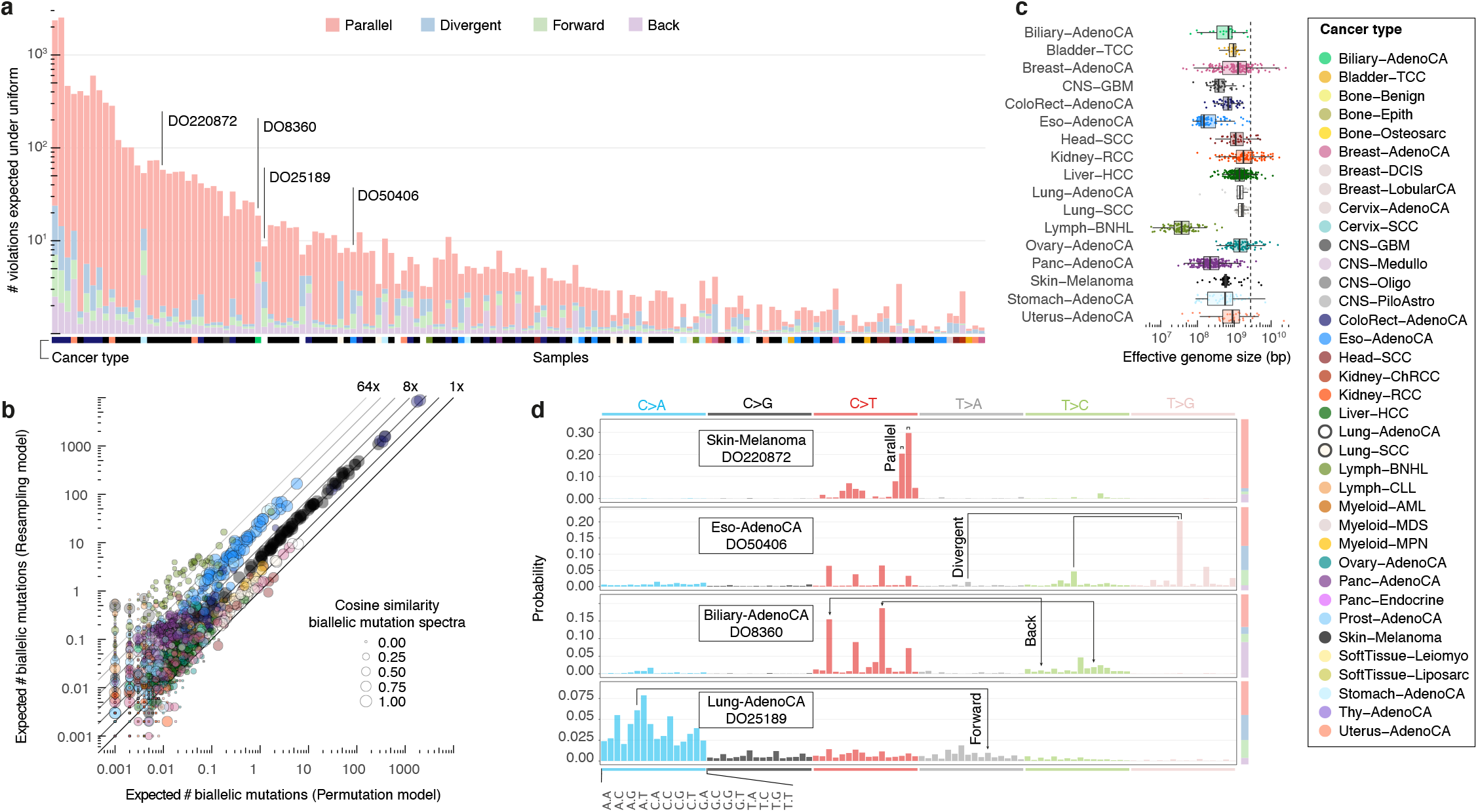
Simulated landscape of infinite sites violations in the PCAWG cohort. (**a**) Number and type of infinite sites violations in 147 PCAWG samples with ≥ 1 expected violation under a uniform mutation distribution. Bar height indicates the expected number of violations and coloured subdivisions represent the fractions contributed by each violation type. Tumour histology of the samples is colour-coded below the bars. The four samples highlighted in (d) are indicated. (**b**) Comparison of the expected biallelic violations from the uniform permutation and neighbour resampling models. Every dot represents a tumour simulated 1000x with each model. Colour and size reflect, respectively, tumour type and the cosine similarity of the predicted infinite sites violation mutation spectra. (**c**) Box and scatterplot showing the effective genome size perceived by the mutational processes per cancer type, as estimated from the per sample differences between simulation approaches. The dashed line indicates the callable genome size. (**d**) Mutational spectra of four tumours with distinct violation contributions indicated in (a). The 16 distinct trinucleotide contexts are provided on the x-axis for C>A type substitutions and are the same for each coloured block. The proportion of parallel, divergent, back and forward mutation is indicated in the stacked bar on the right. Frequent combinations of mutations leading to specific infinite site violations are highlighted.

The distinct preferences for parallel, divergent, forward and back mutation may be understood from the active mutational processes (**Figure 2d**). For instance, the dominant mutagenic activity of UV light in cutaneous melanoma (single base substitution signature 7a/b, SBS7a/b) yields almost uniquely C>T substitutions in CC and CT contexts^10,11^, which can only result in the accumulation of biallelic parallel mutations. In contrast, in a case of oesophageal adenocarcinoma, the interplay between SBS17a and b^10,11^ results in various substitutions of T in a C**<u>T</u>**T context, generating both parallel and divergent variants. Back and forward mutation may occur when the variant allele retains considerable mutability. A case of biliary adenocarcinoma shows SBS1 (ageing) in combination with SBS21 and 44 (mismatch repair)^10,11^, which can result in A[T<>C]G and G[T<>C]G back mutation. An example of forward mutation comes from lung adenocarcinoma, in which tobacco smoking (SBS4)^10,11^ drives C[C>A]C substitutions which can be followed by G[T>A]G, appearing as a single C[C>T]C change.

Encouraged by the simulation results, we set out to directly detect biallelic mutations in the bulk sequenced PCAWG tumours. Parallel mutation increases the variant allele frequency (VAF) and may be distinguished from local copy number gains by comparing the VAF to the allele frequencies of neighbouring heterozygous SNPs, taking tumour purity and total copy number into account (**Methods**). Additionally, when proximal to a heterozygous germline variant, read phasing information can also evidence independent mutation of both alleles. This dual-pronged approach is illustrated for melanoma DO220906, where we identify a total of 480 parallel mutations, 74 of which are supported by phasing data (**Figure 3a,b**, **Table S3)**. Leveraging SNV-SNP phasing, we estimate our VAF-based approach shows a median precision of 86.8% and recall of 54.5% (**Methods**). No parallel mutations are called in regions with loss of heterozygosity, as they cannot be distinguished from early mutations followed by copy number gains.

**Figure 3.**
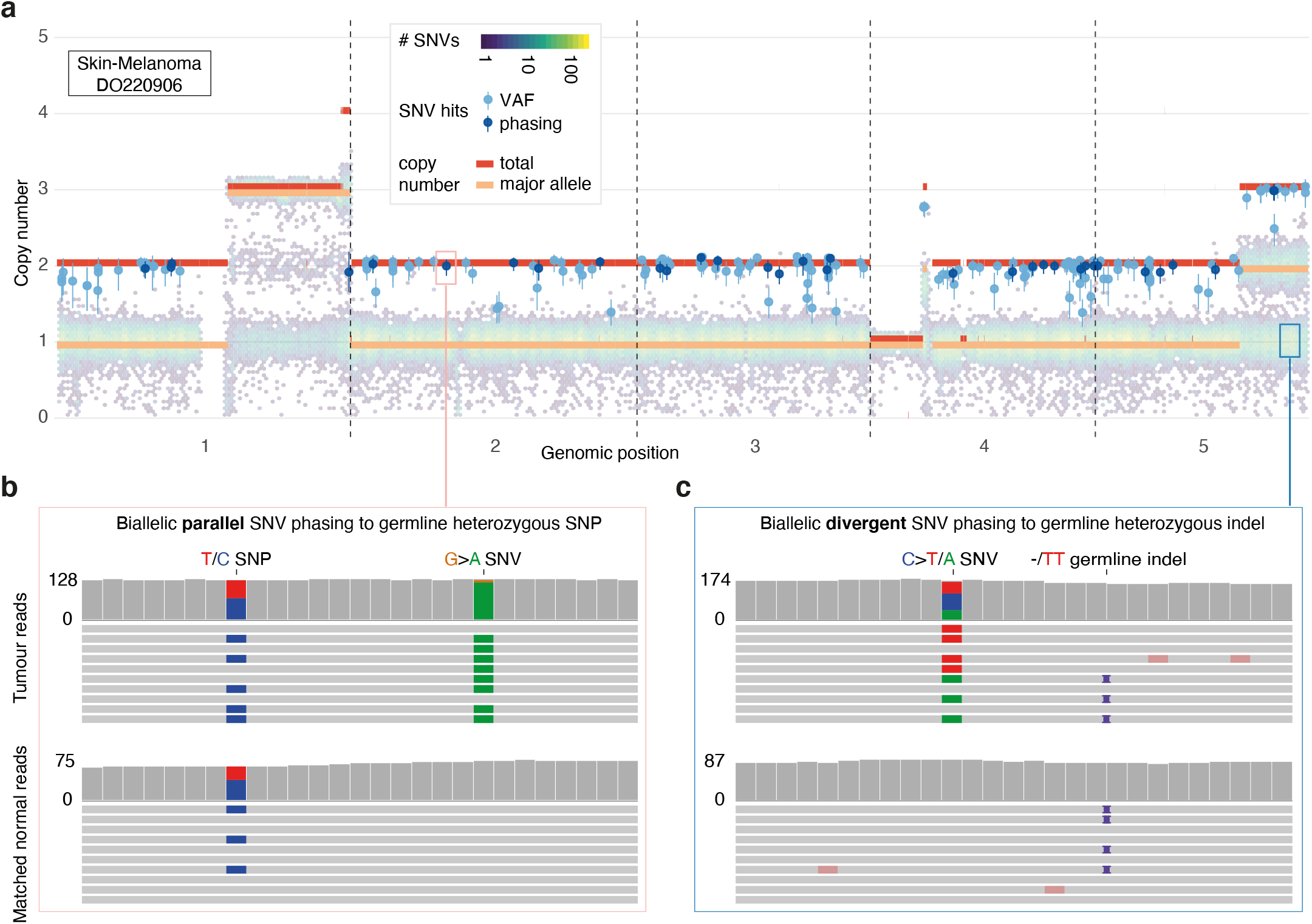
Detecting biallelic mutations in a case of melanoma. (**a**) Tumour allele-specific copy number and binned mutation copy number (hexagons) plotted for chromosomes 1–5 of melanoma DO220906. Somatic SNVs with a mutation copy number exceeding that of the major allele (and equal to the total copy number) are evident, suggesting biallelic parallel mutation events. Error bars represent the posterior 95% highest density intervals. (**b,c**) IGV visualisation of DO220906 tumour (top) and matched normal (bottom) sequencing data at two loci, illustrating how read phasing information can confirm independent mutation of both parental alleles for (**b**) parallel and (**c**) divergent mutations detected after recalling using Mutect2 (**Methods**). Reads (horizontal bars) are downsampled for clarity and local base-wise coverage is indicated left of the histograms.

Likewise, divergent mutations can be picked up by variant callers but are traditionally considered artefacts and filtered out^4^. As neither the PCAWG consensus nor the four contributing variant callers report divergent mutations, we recall mutations with Mutect2 for 195 relevant cases, allowing up to two alternative alleles instead of one (**Methods**). In melanoma DO220906, this yields 8 divergent mutations: 1 with two novel alleles and 7 which add a second alternative allele to a PCAWG consensus variant (**Figure 3c, Table S3–4**). Overall, recalling identifies a median 96.3% of consensus variants and adds 9.5% novel variants, with 0.04% of the latter contributed by divergent mutations (**Figure S1, Table S4**). For 90% of divergent mutations, one of the two alternate alleles is already reported in the PCAWG consensus.

In total, we identify 5,330 divergent mutations, 15,167 parallel SNVs and 29 dinucleotide variants in 722 (26.0%) PCAWG samples (**Tables S4–5**). We reveal 8 candidate biallelic driver events, including parallel nonsense mutations in tumour suppressors *ASXL2* and *CDKN2A* (**Table S6**). VAF outlying parallel mutations confirmed by phasing to proximal SNPs are found in cases of hepatocellular carcinoma and pancreatic adenocarcinoma with as few as 8,892 and 8,941 SNVs (**Figure 4**). Likewise, divergent mutations matching the predicted types are repeatedly identified in oesophageal adenocarcinoma samples with 20,000-30,000 SNVs, while they are absent from melanoma cases with a similar total mutation burden. On the other end of the spectrum, phasing indicates that two ultra-hypermutated colorectal adenocarcinomas each boast around 8,000 parallel and 1,700 divergent mutations.

**Figure 4.**
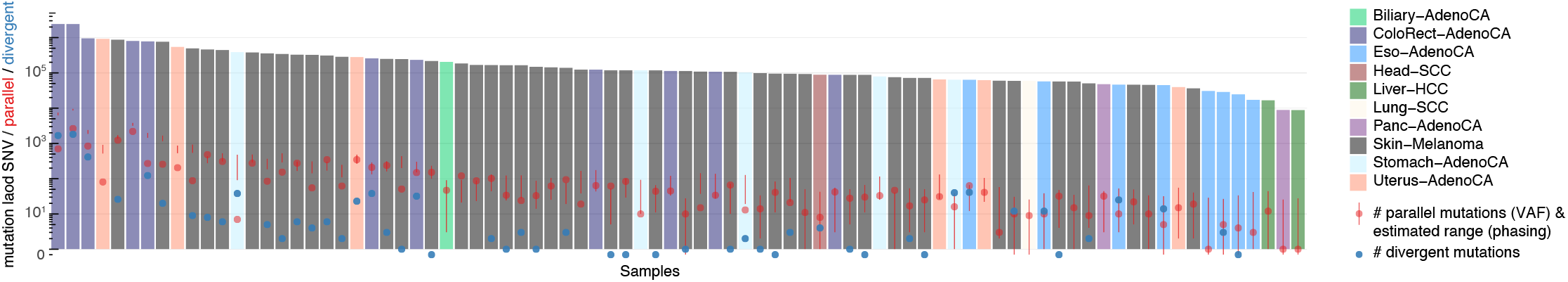
Landscape of biallelic mutations across PCAWG. Number of observed parallel (red) and divergent (blue) mutations plotted in context of the total SNV burden for 84 PCAWG samples with ≥ 1 phasing-confirmed VAF hit. The range of parallel mutations expected purely from SNV-SNP phasing is also indicated (95% confidence interval, red vertical bars) as this approach is less sensitive to purity and copy number state than the VAF-based analysis. Samples for which the number of divergent mutations is not shown, were not considered for Mutect2 recalling.

As hinted above (**Figure 2d)**, biallelic mutations are expected to carry a mutational footprint determined by, but distinct from, the overall mutational profile. For example, as parallel mutations require two independent and identical hits, they are predicted to show a mutation spectrum similar to the square of that of the regular SNVs (**Figure 5a,b**). Indeed, the observed biallelic mutations are better explained by the simulated violation spectra than the overall mutation spectra (*p* = 1.47×10^−2^ and 1.35×10^−8^ for parallel and divergent, respectively, median simulated–observed cosine similarities 0.945 and 0.944, Mann–Whitney *U,* samples with ≥ 10 violations). This further supports the accuracy of our biallelic mutation calls, excluding major contributions from sequencing and alignment artefacts, missed germline variants, undetected focal tandem duplicator phenotypes, precursor lesions or an as yet unknown somatic gene conversion process.

**Figure 5.**
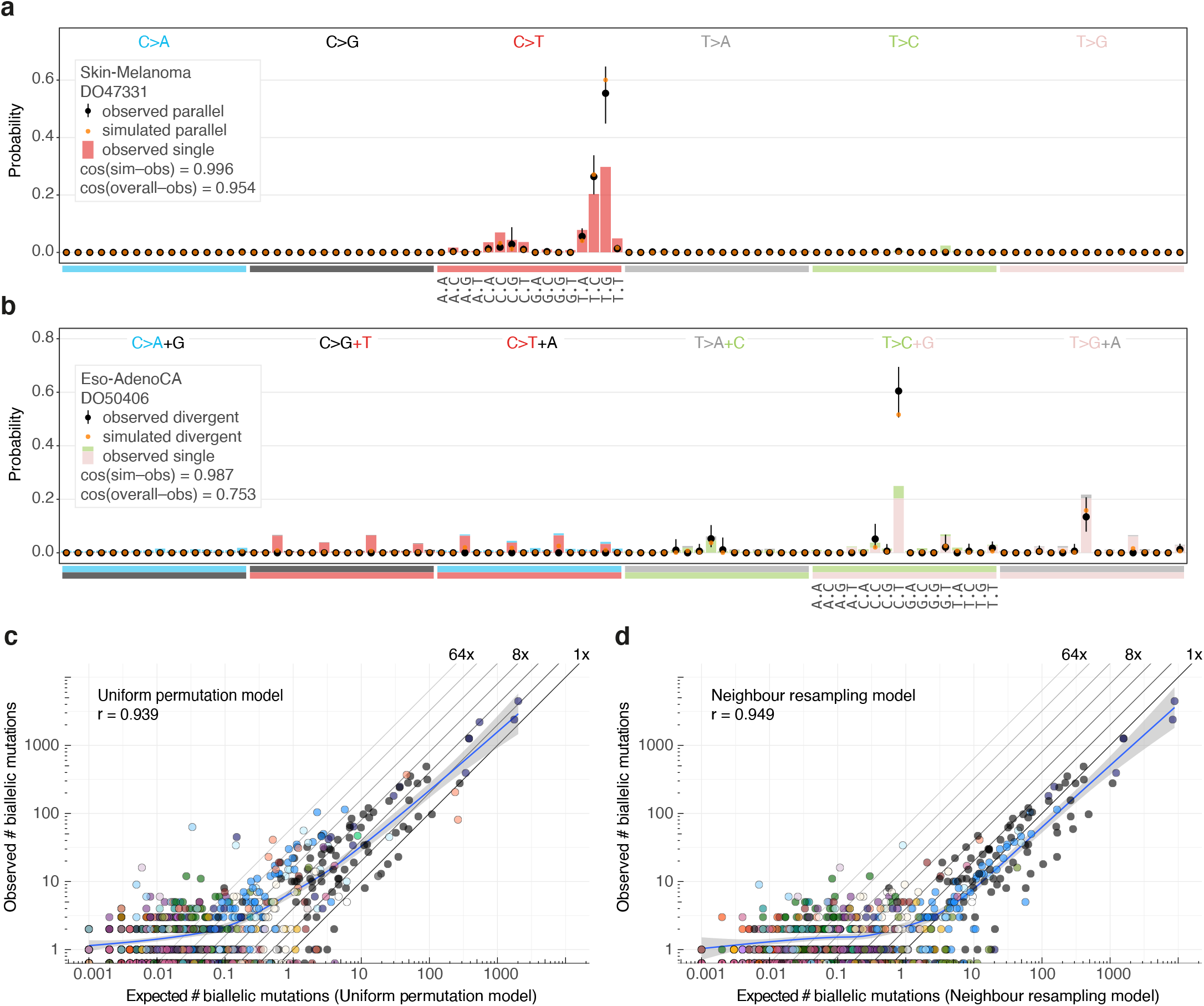
Comparison between observed and simulated biallelic mutations. (**a**) Bar chart highlighting the mutation spectrum of observed and predicted parallel mutations as well as the background SNVs for melanoma DO47331. Cosine similarities between the spectra are indicated. (**b**) Similar as (a) but showing divergent mutations for oesophageal adenocarcinoma DO50406. Bars are stacked to reflect the frequency of the colour-coded base changes indicated on top. Error bars represent the posterior 95% highest density intervals. (**c,d**) Scatterplots of the observed *vs.* expected number of biallelic mutations (parallel + divergent) for all PCAWG samples for the uniform permutation (**c**) and neighbour resampling models (**d**). A spline regression fit is shown together with the Pearson correlation.

While there is a close match between the simulated and observed biallelic mutation spectra, the assumption of a uniform distribution results in a gross underestimate of the number of observable violations. Various melanomas and oesophageal adenocarcinomas harbour over 8 to 32-fold excess biallelic mutations (**Figure 5c**). In contrast, the neighbour resampling model is more accurate, confirming that the effective genome size perceived by various mutational processes is only a fraction of the callable human genome (**Figures 3c** and **5d, Methods**).

Non-uniformity of the mutation rate should result in more biallelic mutations at loci with a higher mutability (*i.e.* hot spots). Indeed, the fraction of loci with biallelic hits can be seen to increase along with the mutation rate as observed in the PCAWG cohort (**Figure S2**). In addition, recurrent biallelic events suggest a high base-wise mutation rate at some positions in the genome (**Figure 6a**). The most frequently hit locus is the promoter of *RPL18A* (chr19:17,970,682), showing three parallel violations and one biallelic variant, accounting for 8 independent hits, all in melanoma (**Figure S3**). Across PCAWG, 9 more melanomas carry monoallelic SNVs at this position (12% total)^13^. Differential motif enrichment at loci with biallelic *vs.* trinucleotide-matched monoallelic hits in melanoma reveals enrichment of Y**<u>C</u>**TT**<u>C</u>**CGG and WTTT**<u>C</u>**C motifs (**Figure 6a,b**)^14^. Y**<u>C</u>**TT**<u>C</u>**CGG motifs are recognised by E26 transformation-specific (ETS) transcription factor family members. Binding has been shown to render them more susceptible to UV damage due to a perturbation of the Tp**<u>C</u>** C5– C6 interbond distance *d* and torsion angle *η*, favouring cyclobutane pyrimidine dimer formation (**Figure 6c,d**)^15,16^. The WTTT**<u>C</u>**C motif matches the recognition sequence for Nuclear factor of activated T-cells (NFAT) transcription factors^17,18^. Analysis of crystal structures of NFATc1–4 in complex with its cognate DNA indicates that binding induces similar, albeit less outspoken, conformational changes to the Tp**<u>C</u>** dinucleotide which may explain its increased UV-mutability (**Figure 6d**).

**Figure 6.**
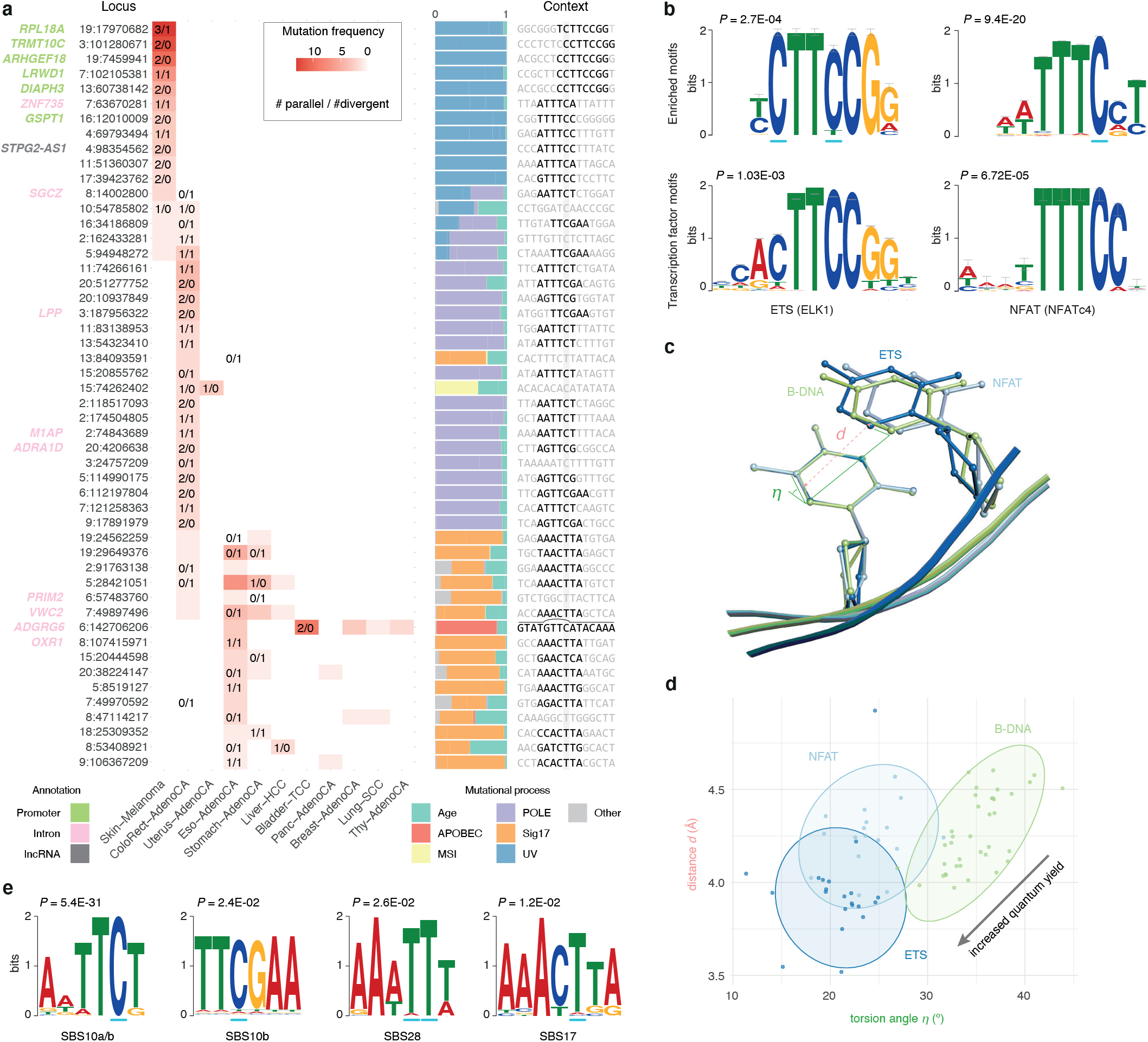
Biallelic mutations reveal tumour type-specific mutational hot spot contexts. (**a**) Heatmap of the fifty most frequently mutated loci in PCAWG with at least one biallelic mutation. The number of parallel/divergent mutations at each site is indicated, as are gene annotations, the underlying mutational processes, and the local sequence context with emerging motifs. For chr6:142,706,206, part of the stem and loop of a local sequence palindrome are indicated. MSI, miscrosatellite instability. (**b**) Sequence logos of motifs enriched at loci with biallelic mutations in melanoma (top) and corresponding transcription factor recognition sequences (bottom). (**c**) Superposition of TpC dinucleotides in crystal structures of ETS-bound (GABP), NFAT-bound (NFAT1c) and free B-DNA (PDB IDs, 1AWC, 1OWR and 1BNA, respectively). The distance *d* between the midpoints of the two adjacent C5–C6 bonds as well as their torsion angle is indicated. (**d**) Scatter plot showing the distances and angles indicated in (c) as observed in crystal structures from the RCSB protein data bank. (**e**) Sequence logos of motifs enriched at loci with biallelic mutations in colorectal adenocarcinoma (SBS10, 28) and oesophageal/stomach adenocarcinoma (SBS17).

Motif enrichment analysis on bi- *vs*. monoallelic sites from colorectal adenocarcinoma reveals special cases of the sequence contexts of SBS10a/b and SBS28, which are associated with Pol *ε* exonuclease domain mutations (**Figure 6a,e**)^10,11,19^. AWTT**C**T and TT**C**GAA carry extra adenosine and thymine bases surrounding the regular trinucleotide context of the mutated C in SBS10, a preference also observed in the recent extension from tri- to pentanucleotide contexts^11^. It is unclear how these additional bases contribute to the mutability of these motifs. A 5mC mutator phenotype of *POLE-mutant* cancers has been described however^20^, and we confirm methylation of these loci in normal colon (median methylation rate 0.84, one-sided Mann–Whitney *U* test vs. background, *p* = 4.87×10^−5^), providing context for the latter motif. In case of the SBS28 hypermutable motif AAA**<u>TTT</u>**, the presence of an AAA stretch upstream of the mutated T is also yet to be explained. Likewise, AT-rich sequences surrounding the canonical SBS17-mutable trinucleotide context C**<u>T</u>**T can render some loci hypermutable in oesophageal and stomach adenocarcinomas (AAAC**<u>T</u>**TA motif; **Figure 6a,e**). Pentanucleotide mutational signatures confirm the local AT-bias^11^ and it is tempting to speculate secondary structure could be involved.

Last, it is worth highlighting recurrent (biallelic) mutation at chr6:142,706,206, in an intron of *ADGRG6* (**Figure 6a**). The CTCTTTGTAT-GTT**<u>C</u>**-ATACAAAGAG palindromic sequence may adopt a hairpin structure, exposing the hypermutable C at the last position of a 4bp loop and rendering it susceptible to APOBEC3A deamination, in line with recent findings^7^.

Taken together, we identify 20,555 biallelic mutations in 26% of PCAWG cases, demonstrating how the infinite sites model breaks down at the bulk level for a considerable fraction of tumours. By extension, the model is untenable in most, if not all, tumours at the multi-sample or single cell level, as violations become increasingly frequent for larger sets of mutations and lineages (**Figure S4**). If not correctly identified, biallelic mutations confound variant interpretation, ranging from driver inference to subclonal clustering and timing analyses, as well as phylogenetic inference. Nevertheless, at-scale detection of biallelic mutations affords an intimate look at previously hidden features of the mutational processes operative in cells, such as hot spots, hypermutable motifs and the molecular mechanisms of DNA damage and repair. These observations underscore the need for accurate models of mutation rates and tumour evolution as well as careful interpretation of allele frequencies, phasing data and driver inference.

## Supplementary figure legends

**Figure S1.**
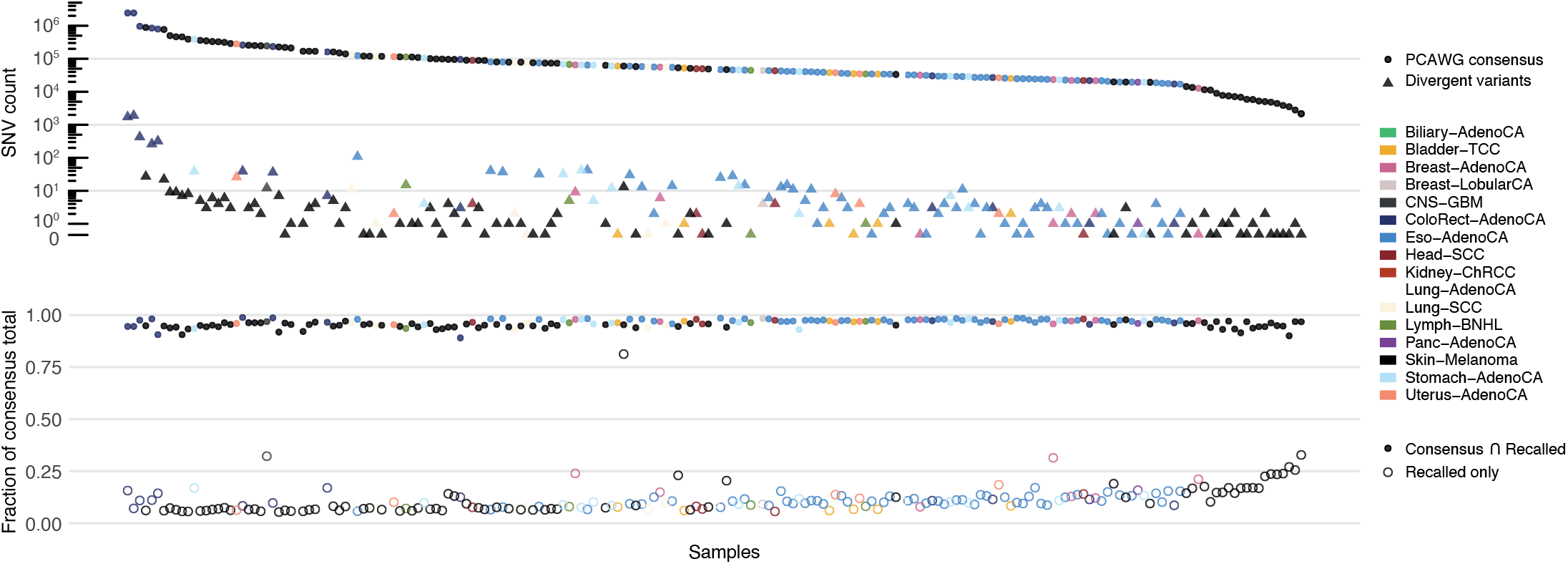
Variant recalling results on 195 PCAWG tumours. Dot plot showing the total number of PCAWG consensus SNV calls and the number of divergent mutations identified after recalling with Mutect2 (top). Fraction of PCAWG consensus calls recovered during recalling and fraction of new calls (bottom).

**Figure S2.**
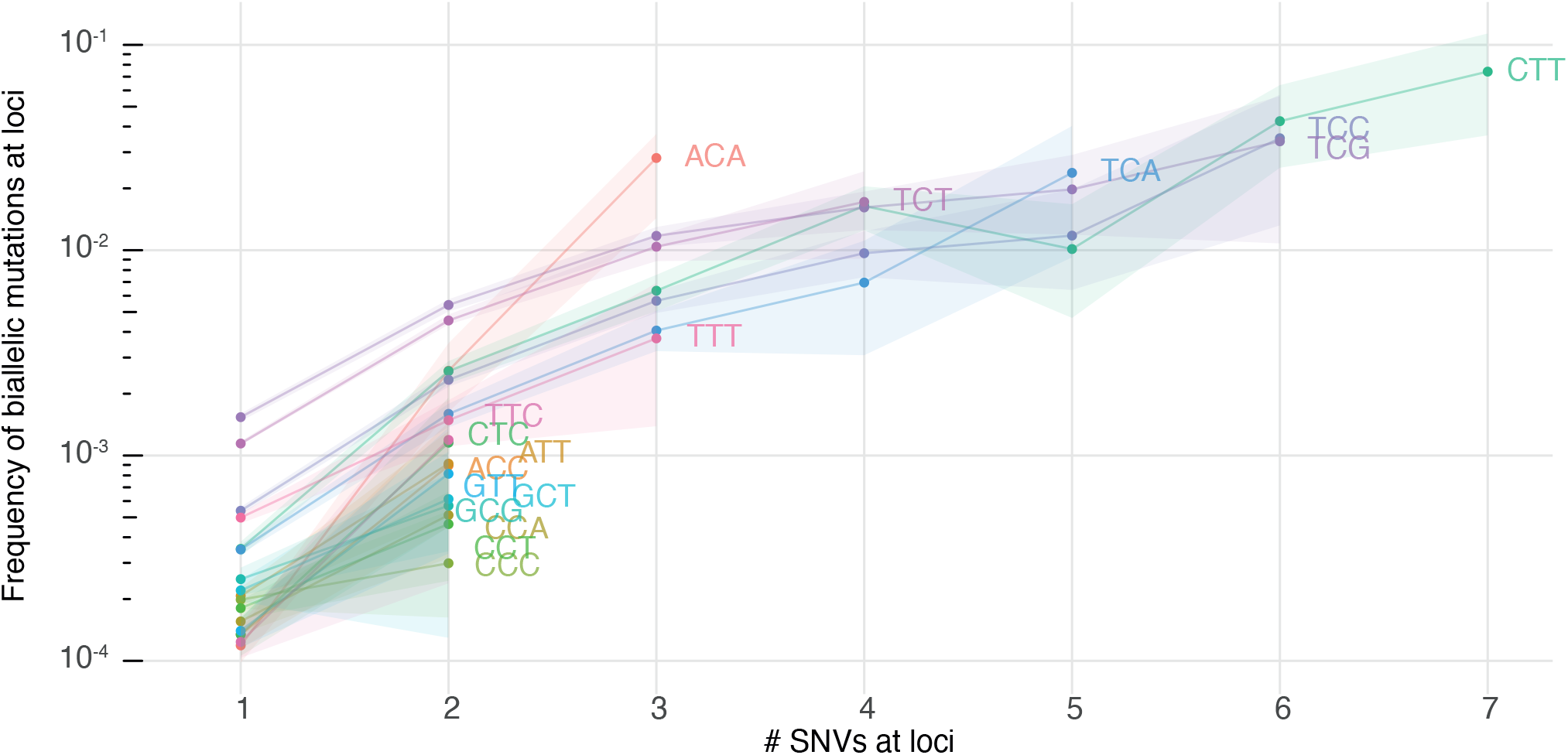
Loci with biallelic mutations have higher intrinsic mutability. The fraction of loci with biallelic mutations is plotted for loci with 1, 2,…,7 monoallelic SNVs across PCAWG. Loci are further stratified per trinucleotide context. Bootstrap resampling is performed to obtain 95% confidence intervals.

**Figure S3.**
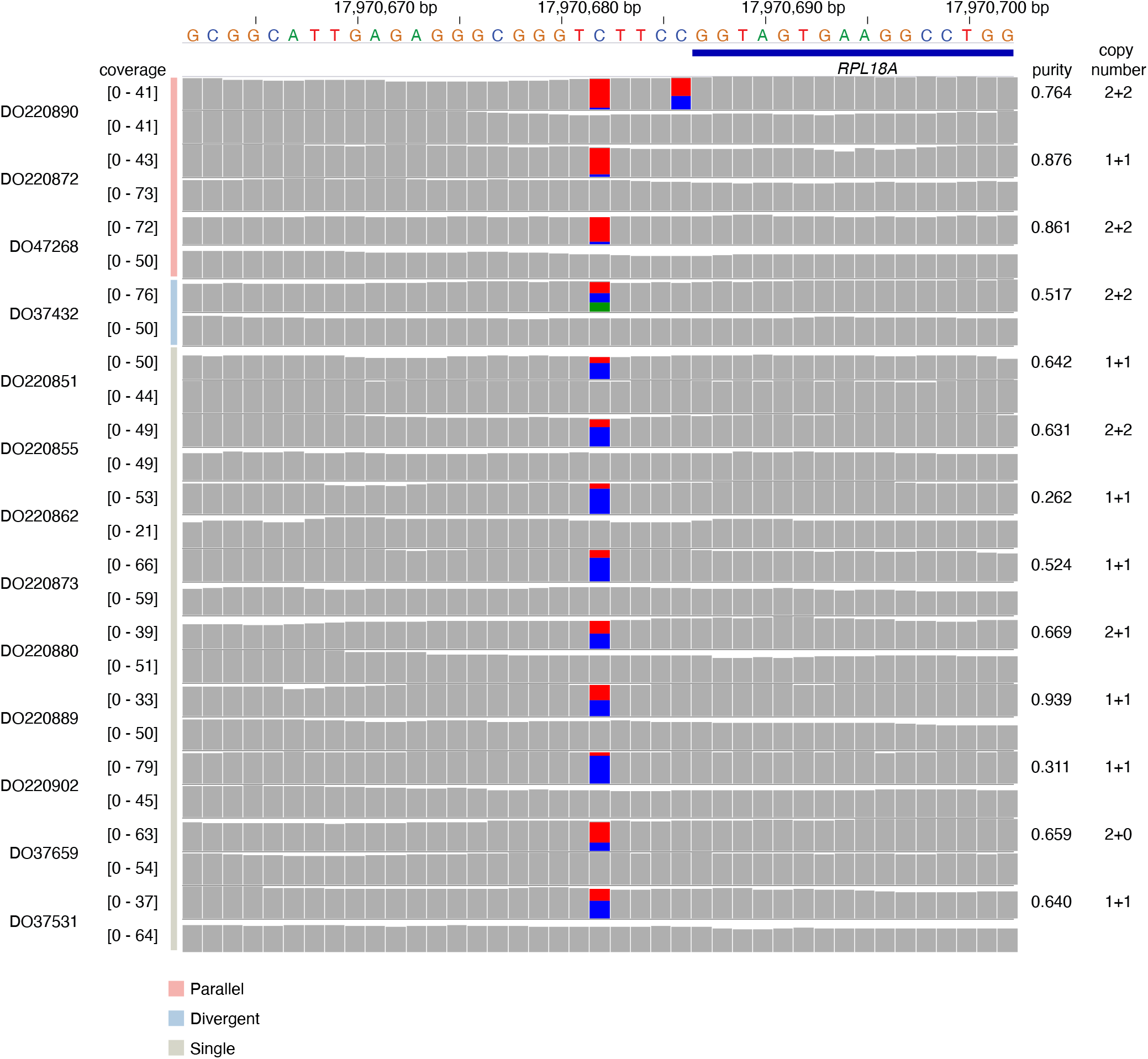
Recurrent mono- and biallelic mutation of the *RPL18A* promoter. Histograms of read and base coverage in 13 melanoma tumour-normal pairs showing mono- or biallelic mutation of the ETS-binding T**<u>C</u>**TTCCG motif at the *RPL18A* promoter.

**Figure S4.**
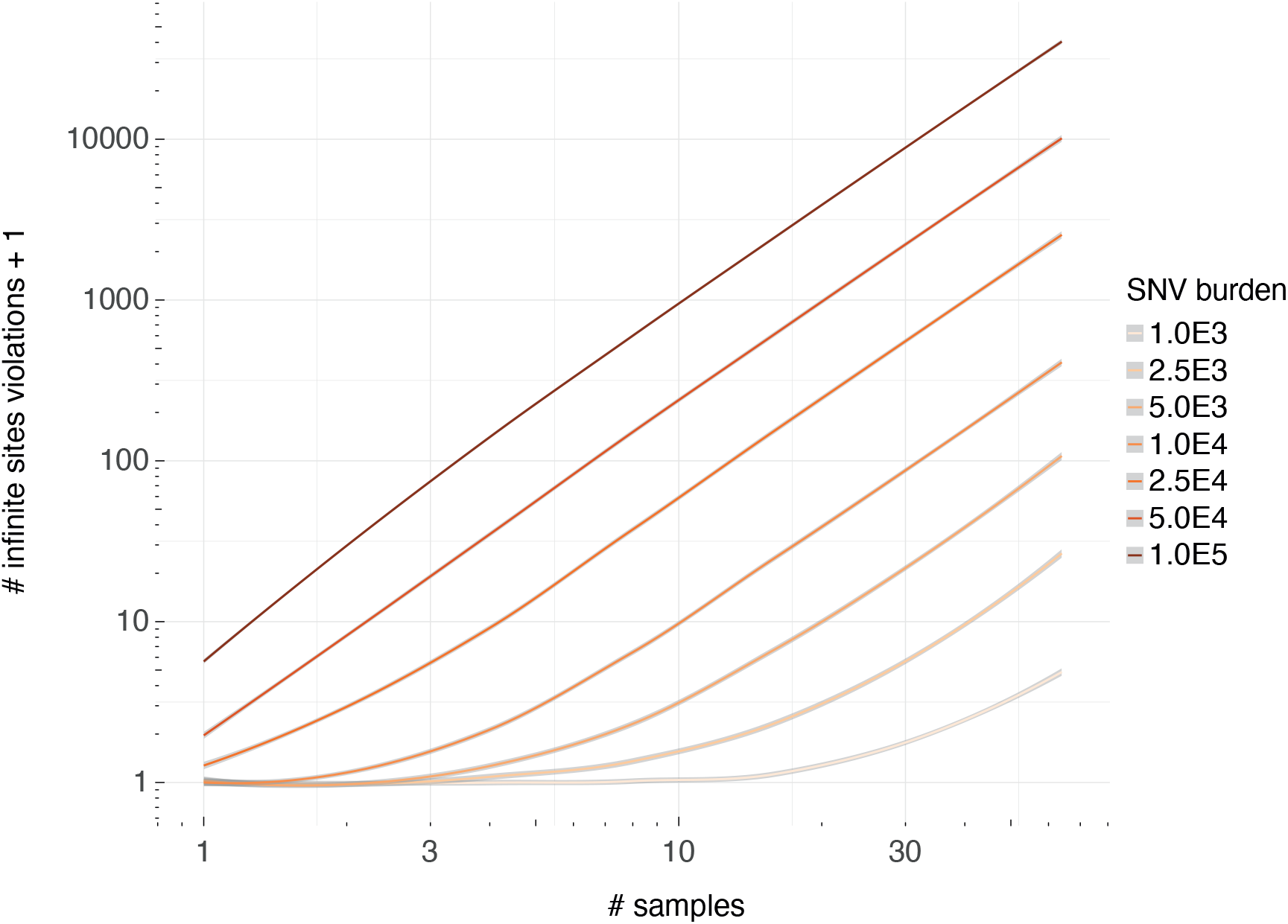
Infinite sites violations in a multi-sample setting. Simulation results showing how the number of infinite sites violations increases when multiple samples are considered, each with the indicated mutational load (coloured lines). Gray bands indicate 95% confidence intervals of a spline fit.

## Supplementary tables

**Table S1 | Uniform permutation-based infinite sites simulation results**

Number of biallelic parallel and divergent, and forward and backwards-type infinite sites violations in 1000 simulations using a uniform permutation approach across the callable genome

**Table S2 | Neighbour resampling-based infinite sites simulation results**

Number of biallelic parallel and divergent-type infinite sites violations in 1000 simulations of a resampling-based approach using tumours of the same cancer type with similar mutational processes.

**Table S3 | Mutect2 variant calling in 195 PCAWG samples**

For each sample – selected for its likelihood of harbouring biallelic divergent mutations or belonging to the same cohort of samples with a high likelihood – variant calls are compared to the PCAWG consensus SNV calls and the number of biallelic mutations is given.

**Table S4 | Biallelic divergent mutations in 195 PCAWG samples**

List of identified biallelic divergent mutations with read counts and additional quality control metrics.

**Table S5 | Biallelic parallel mutations in PCAWG**

List of all identified biallelic parallel mutations in PCAWG with read counts and additional quality control metrics.

**Table S6 | Candidate biallelic driver mutations**

List of all nonsynonymous biallelic mutations in known cancer driver genes (COSMIC and PCAWG consensus driver gene lists).

## Acknowledgements

This work was supported by the Francis Crick Institute, which receives its core funding from Cancer Research UK (FC001202), the UK Medical Research Council (FC001202), and the Wellcome Trust (FC001202). This project was enabled through access to the MRC eMedLab Medical Bioinformatics infrastructure, supported by the Medical Research Council (grant number MR/L016311/1). JD is a postdoctoral fellow of the European Union’s Horizon 2020 research programme (Marie Skłodowska-Curie Grant Agreement No. 703594-DECODE) and the Research Foundation–Flanders (FWO 12J6916N). PVL is a Winton Group Leader in recognition of the Winton Charitable Foundation’s support towards the establishment of The Francis Crick Institute. This research was funded in part by the Wellcome Trust (FC001202). For the purpose of Open Access, the authors have applied a CC BY public copyright licence to any Author Accepted Manuscript version arising from this submission. The authors would like to thank Paul C. Boutros for constructive criticism of the manuscript.

## Online methods

### Singe Nucleotide Variant calling

Consensus single and multi-nucleotide variant calls are obtained from the ICGC-TCGA PCAWG project^12^. Briefly, these calls were constructed according to a “2+ out of 4” strategy, where calls made by at least two callers (the three Broad, EMBL/DKFZ, and Sanger core PCAWG pipelines, plus MuSE v1.0) were selected as consensus calls. Post-merging, these calls were subject to further quality control including filtering against oxidative artefacts (OxoG) and alignment (BWA *vs.* BLAT) or strand biases resulting from different artefact-causing processes, as well as checks for tumour-in-normal and sample cross-contamination. Crucially, great care was taken to avoid “bleed-through” of germline variants into the somatic mutation calls. Specifically, absence from the Broad panel-of-normals based on 2,450 PCAWG samples and a higher read coverage (≥19 reads with at most one read reporting the alternate allele) in the matched normal sample were required to call a somatic mutation at one of the >14M common (>1%) polymorphic loci of the 1000 genomes project. SNVs that overlapped a germline SNV or indel call in the matched normal were also removed. Sensitivity and precision of the final consensus somatic SNV calls were 95% (90% confidence interval, 88–98%) and 95% (90% confidence interval, 71–99%), respectively, as evaluated by targeted deep-sequencing validation^12^. Of note, 18 biallelic parallel mutations identified here were covered by the PCAWG validation effort with 17 passing and one not being observed.

To identify biallelic divergent variants, which are filtered out in PCAWG, we recalled variants on 195 non-graylisted PCAWG tumour-normal pairs (that do not show any tumour-in-normal contamination) where we might reasonably expect to find such mutations according to our uniform permutation simulations. Included also, as an internal control, are all other samples from the Australian PCAWG melanoma cohort (MELA-AU) which meet these criteria but in which we do not expect biallelic divergent mutations. SNVs and indels are called using Mutect2 (GATK v4.0.8.1) on the base quality score-recalibrated PCAWG bam files and filtered following best practices^21^. The Genome Aggregation Database (gnomAD) was provided as a germline resource for population allele frequencies and an additional panel of normals was also derived from all matched normal cases. To prevent filtering of biallelic variants, FilterMutectCalls is run with the --max-alt-allele-count flag set to 2. Additional filtering against potentially missed germline SNPs was done by requiring a posterior probability for the alternative allele to be germline (P_GERMLINE) < −1 for both of the alternate alleles and requiring a minimal depth of 19 high quality reads (mapping quality ≥ 35 and base quality ≥ 20) in the matched normal sample.

### Consensus copy number, purity and ploidy

PCAWG consensus copy number profiles were obtained from Dentro *et al.*^3^. Briefly, we first segmented each cancer’s genome into regions of constant copy number using six individual copy number callers. Segment breakpoints were based on the PCAWG consensus structural variants (SVs) complemented with high-confidence breakpoints identified by several of the copy number callers. The six callers were then re-run, enforcing this consensus segmentation as well as separately established consensus tumour purity and ploidy values, to determine the allele-specific copy number of each segment. The allele-specific copy number calls were then combined into a consensus profile using a multi-tiered approach and segments were assigned a level of confidence.

### Simulating infinite sites violations

To estimate the number of infinite sites violations in tumours, we developed two distinct simulation approaches leveraging the SNV calls in the PCAWG cohort.

Our first simulator (termed uniform permutation model) resamples the observed SNVs in a tumour uniformly across the callable regions of the chromosomes, according to the observed trinucleotide-based mutational spectrum. A single simulation proceeds as follows. First, the total mutational load *n_t,sim_* is resampled from a gamma–Poisson mixture where the Poisson rate parameter *λ* ~ Gamma with mode equal to the observed mutational load *n_t,obs_* and a standard deviation *σ* = 0.05 × *n_t,obs_*. That is: *n_t,sim_* ~ *Poisson*(*λ* ~ *Gamma*(*r,β*)) where the rate of the Gamma distribution 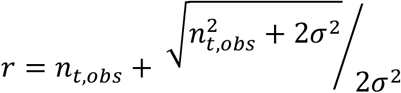 and the shape *β* = 1 + *n_t,obs_* × *r*. Mimicking the observed distribution, these mutations are then divided across the chromosomes according to a Dirichlet-multinomial model with *n_t,sim_* trials and parameter vector ***α*** where *α_i_* is equal to 1 + the total mutational burden on chromosome *i*. That is: ***n_sim_*** ~ *Mult* (*n_t,sim_*, *π* ~ *Dir*(***α***)) with ***α*** = (*n*_1,*obs*_, *n*_2,*obs*_,..., *n_X,obs_*) + **1**. Next, mutation spectra per chromosome (***π_i_***) are sampled from a Dirichlet distribution with parameter vector ***μ_i_*** where *μ_i,j_* is equal to a pseudocount *ψ_j_* derived from the overall mutational spectrum plus the observed number of mutations of type *j* on chromosome *i*. That is: ***π_i_*** ~ *Dir*(***μ_i_***) with 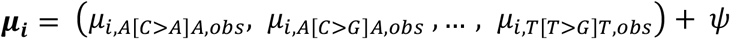 with 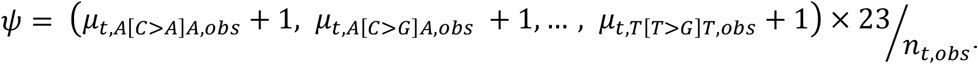 These spectra are then normalised to mutation type probabilities using the trinucleotide content on the corresponding chromosomes. In turn, the probabilities are used for rejection sampling of *n_i,sim_* mutations at trinucleotides sampled uniformly along the two (diploid) copies of the callable parts of chromosome *i*. The resulting mutation spectra are indistinguishable from the observed spectrum of the sample. During simulation, the algorithm keeps track of which allelic positions have been mutated and considers them accordingly for (i) biallelic parallel mutation, where two alleles independently mutate to the same alternate base; (ii) biallelic divergent mutation, by independent mutation of two alleles each to another base; (iii) back mutation, whereby an earlier introduced variant is mutated back to the wild type; and (iv) forward mutation, where an earlier variant is mutated to another. Simulations are repeated 1,000 times for each sample and the totals, median and 95% intervals are reported for each violation type and context.

In a second simulation approach (termed neighbour resampling model), we resample without replacement the mutational landscape of a tumour from the pooled SNVs of representative PCAWG tumours. In this context, we define a tumour as representative for the simulation target when it is of the same tumour type (PCAWG histology) and has similar mutational signature exposures (cosine similarity of their mutation spectra ≥ 0.9). Note that this approach allows to simulate biallelic events but not back and forward mutation. We further exclude all graylisted and non-preferred multi-sample tumours as well as 21 prostate cancer cases from the PRAD-CA cohort which were suspect of contamination harbouring excess low VAF SNV calls in repetitive regions.

### Identification of parallel mutations – allele frequencies

Parallel mutation increases the variant allele frequency, which can be picked up by comparing it to the B-allele frequency (*BAF*) of local heterozygous SNPs, taking tumour purity and local total copy number (log R) into account. In the first part of the approach, we obtain phased *BAF* values and log R as an intermediate output of the Battenberg copy number calling pipeline^3^. Briefly, allele counts at 1000 Genomes v3 SNP loci are extracted from the matched tumour and normal bam files using alleleCount with a minimal base quality of 20 and mapping quality of 35. Heterozygous SNPs are identified as having 0.1 < *BAF* < 0.9. in the matched normal sample and poorly behaving loci are filtered out (Battenberg problematic loci file). Haplotypes are imputed using Beagle5 followed by a piecewise constant fit of the phased tumour *BAF* values and flipping of haplotype blocks with mean *BAF* < 0.5. Total allele counts of tumour and normal are converted into Log R values and corrected for GC-content and replication timing artefacts.

*BAF_seg_* and *log R_seg_* estimates are computed for all PCAWG consensus copy number segments^3^. Allele counts at phased heterozygous SNPs are considered to be generated according to a beta-binomial model with *V_i_* ~ *Bin*(*n_i_* = *V_i_* + *R_i_,p* ~ *Beta*(*BAF_seg_* × *ψ*,(1 – *BAF_seg_*) × *ψ*)) where *V_i_* and *R_i_* are, respectively, the observed counts of the major and minor allele of SNP *i*, and *ψ* is a sample-specific concentration parameter (*i.e.* a pseudo-coverage of the average segment). For each sample, *ψ* is optimised between 50 and 1000, by computing for each SNP a two-sided *P*-value from the beta-binomial model above and ensuring the robustly fitted slope of a QQ-plot of these *P*-values is equal to 1.

A similar model can subsequently be used to test whether a variant is present on a higher number of copies than the number of copies of the major allele present in the tumour. In pure tumour samples, this would be directly observable as their allele frequency exceeds that of local heterozygous SNPs on the major allele. Considering admixed normal cells, however, the maximal expected allele frequency needs to be corrected for tumour purity and total copy number of the segment. This corrected “somatic” BAF can be derived as follows:

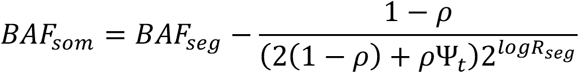

with *ρ* and Ψ_*t*_, the PCAWG consensus tumour purity and ploidy (*i.e.* the average tumour copy number), respectively^3^. This amounts to subtracting from the segment *BAF* the contribution of the major allele from admixed normal cells.

The final beta-binomial model with *BAF_som_* and *ψ* then describes the expected allele counts of clonal somatic variants carried on all copies of the major allele. This model is used to perform independent filtering and assess powered loci using a one-sided test for the SNVs contained on that copy number segment as *P*(*V_i_* ≥ *v* | *V_i_* + *R_i_*, *BAF_som_,ψ*). *P*-values are corrected for multiple testing according to Benjamini–Hochberg and SNVs are considered as potential parallel mutations when *q* ≤ 0.1.

A number of additional quality checks and filters are in place to mitigate effects of potential errors and biases in allele counts, consensus genome segmentation, purity and ploidy:

i. SNVs overlapping a known (1000 genomes v3) heterozygous germline SNP in the individual are filtered out.
ii. The robustly fitted slope of a QQ-plot of the final SNV *P*-values should be ≤ 1, if not, sample purity may have been underestimated and the sample is excluded.
iii. Candidate parallel mutations with ≥ 2 heterozygous SNPs within 25 bp are filtered out as these affect mapping qualities and bias allele counts.
iv. SNVs in regions with inferred loss of heterozygosity (copy number of the minor allele equal to 0) in the PCAWG consensus copy number are not tested. Similarly, in males, only the pseudoautosomal regions of X are considered.
v. *BAF* and *log R* of proximal heterozygous SNPs on either side of a candidate variant should not represent outliers on the segment as a whole, which could indicate a missed copy number event. For the *BAF,* we require the two-sided beta-binomial *P*-values of these SNPs, as computed above, to be > 0.001 and their combined *P*-value (Fisher’s method) > 0.01. For the *log R,* identical thresholds apply, with *P*-values derived using a two-tailed test assuming a Gaussian distribution with mean equal to the median segment *log R* and standard deviation the median absolute deviation adjusted for asymptotic consistency.
vi. If *BAF_som_* is estimated to be < 0.05 for a segment, it is conservatively raised back to *BAF_seg_*.
vii. Candidate variants from tumours in which neither the permutation nor the resampling-based simulator yielded any biallelic mutations across 1000 simulations were excluded.

Further flags were included for quality control, but were not used during filtering of the final call set. (i) Candidate biallelic hits at T- and B-cell receptor loci are flagged to assess the impact of small V(D)J recombination-derived deletions in infiltrating immune cells on allele frequencies and coverage. (ii) For each variant, we checked whether it lifted over from the 1000 Genomes GRCh37 build (hg19) to a single location on the hg38 assembly, also requiring the same reference base at that position. (iii) SNVs were flagged if near a somatic or germline indel (position −10 to +25) in the same sample.

### Identification of parallel mutations – variant phasing

Phasing information is obtained for all heterozygous SNP–SNV pairs that are within 700bp of one another. We apply the following stringent filtering: count only read pairs with mapping quality ≥ 20, mismatch bases quality ≥ 25, no hard or soft-clipping, that are properly paired, are not flagged as duplicates and do not have a failed vendor quality control flag. Furthermore, we remove read pairs with indels and those that have ≥ 2 mismatches in a single read or ≥ 3 in the whole pair (if both phased variants are spanned by different reads in the pair).

We infer a parallel mutation when, for a heterozygous SNP–SNV pair, at least 2 reads from each allele of the SNP report the somatic variant, *i.e.* at least 2 Ref-Alt and 2 Alt-Alt reads. In addition, Ref-Alt and Alt-Alt reads each should represent > 10% of the total phased reads. To avoid a scenario where, after a gain of the chromosome copy carrying the somatic variant, the phased allele of the heterozygous SNP is mutated to the non-phased allele, we require that the BAF of this SNP is not an outlier on the segment. As described above, this is accomplished by demanding that its two-sided beta-binomial *P*-value > 0.001.

While phasing info is sparse, it is less dependent on the local copy number state, purity and coverage than the VAF approach detailed above. For instance, in contrast to tests on the allele frequency, phasing to a heterozygous SNP can detect parallel mutations on a segment with copy number 2+1 where both parental alleles have only one copy mutated. Phasing results may therefore be used to evaluate the performance of the VAF approach in a sample. However, both approaches are effectively blind in regions with loss of heterozygosity. Parallel mutations can occur in these contexts when the copy number ≥ 2 but cannot readily be distinguished from early mutations which have occurred before the duplication.

Precision and recall of the VAF-based approach are assessed by taking all phaseable SNVs (*i.e.* SNP-SNV pairs having ≥ 2 reads each for the SNP Ref and Alt alleles and ≥ 4 reads reporting the somatic variant) which have been evaluated in the VAF pipeline. Precision is calculated as the fraction of VAF-inferred biallelic parallel mutations which are confirmed by phasing. Recall is the fraction of phasing hits picked up through their allele frequencies. Overall performance is reported as the median precision and recall for samples with ≥ 10,000 phaseable SNVs.

By extrapolating the rate of parallel mutation at phaseable SNVs to all testable SNVs (*i.e.* those passing the quality checks and filters listed above), we estimate the total number of parallel mutations in a sample *i* (*n_viol,i_*). The estimate and its uncertainty can be described using a beta-binomial: *n_viol,i_* ~ *Bin*(*n* = *n_i_,p* ~ *Beta*(*n_phas,par,i_* + 0.001, *n_phas,single,i_* + 0.001)) where *n_i_* is the total number of passed SNVs, *n_phas,par,i_* is the number of phasing-informed biallelic parallel mutations and *n_phas,single,i_* is the number of phaseable SNVs with no phasing evidence for parallel hits.

### Birthday problem approximation

The total number of infinite sites violations in a sample may also be roughly approximated by a variant of the birthday problem, which asks for the probability that at least two people share a birthday in a group of *N* random people. While this simplification ignores intricacies of genomes such as mutation types and copy number, it provides a reasonable first approximation and straightforward mathematical formulation. We start with the probability that mutation A and B hit the same locus, *i.e.* they violate the infinite sites model 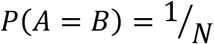 where *N* is the size of the genome. From this it is easy to derive the probability they do not share a locus 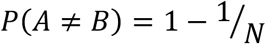. The probability *A* does not hit the same locus as *n* other mutations is then 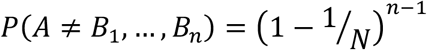. To obtain the expected number of mutations not sharing a locus, this probability is multiplied by the total mutation burden *n*. Finally, the number of infinite sites violations is then 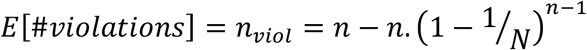. Given that for a human genome 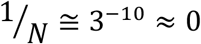, Taylor approximation yields 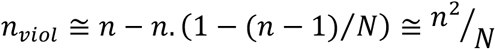, indicating that the number of infinite sites violations scales with the square of the total mutation burden and the inverse of the genome size.

### Motif enrichment

To assess enrichment of specific motifs at sites with biallelic mutations, we extracted 15bp sequence contexts (+ strand where C or T is the reference base and - strand otherwise), for all parallel and divergent biallelic mutations in melanoma, colorectal, oesophageal and stomach adenocarcinomas. For every biallelic mutation, we sampled 10 mutation type-matched (trinucleotide context + alternate base) somatic SNVs from the same tumour and extracted their 15bp contexts as a control set. The Multiple EM for Motif Elicitation suite of tools (STREME and TomTom; v5.3.2) was used to discover sequence motifs enriched in the biallelic set relative to the control set^14^. In the case of melanoma, identified motifs linked to known TF recognition sequences from the HOCOMOCO Human v11 Core collection^18^. *P*-values were computed according to STREME and TomTom.

### Structural analysis

Crystal and NMR-structures for free B-DNA, NFAT- or ETS-bound DNA were obtained from the RCSB Protein Databank. The C5–C6 interbond distances *d* and torsion angles *η* were extracted using PyMOL at the relevant TpC dinucleotide in the ETS and NFAT recognition motifs and at non-terminal TpC dinucleotides in the free B-DNA. When multiple chains were present in a single structure, the average *d* and *η* were used.

